# Mucus-associated microbiota consortia: Insights from the advanced TIM-2*muc* model

**DOI:** 10.1101/2025.07.16.665087

**Authors:** Sjoerd Ruizenaar, Katja Lange, Olaf Heckert, Jan Lelieveld, Andrei Hrynevich, Daniela Pacheco, Lars Verdurmen, Maitrayee Chatterjee

## Abstract

The mucus layer lining the epithelium of the colon contains a microbiome that is distinct from the luminal microbiome; this is referred to as the mucus-associated microbiome (MAM). Recently, research on the MAM has gained momentum due to its significant role in immune and metabolic health. However, the study of the MAM is limited by conventional experimental techniques, as sampling this microbiome often requires invasive procedures, making *in vivo* research challenging. Additionally, obtaining samples from the MAM can compromise the lumen, complicating simultaneous studies of both the MAM and the lumen.

To address these limitations, we present the TIM-2*muc*, an advanced version of the colonic *in vitro* model TIM-2, which includes a mucus compartment comprising both colonic mucus beads - Gut^3^Beads and an electrospun sheet coated with Gut^3^Gel. To validate whether the integrated mucus compartment behaves like the *in vivo* MAM, we examined the bacterial composition, alpha and beta diversity, and the MAM’s composition compared to the lumen compartment of both the TIM-2*muc* and the conventional TIM-2 model.

Our findings suggest that the MAM in the TIM-2*muc* model closely resembles its *in vivo* counterpart, thereby enhancing the physiological relevance of the luminal microbiome. TIM-2*muc*, an advanced TIM-2 model, presents a novel, precise, standardized, and non-invasive method for studying the MAM and LM simultaneously.

## Introduction

The colon houses a plethora of microorganisms in the lumen that together compose the colonic luminal microbiota (LM) (1). The colon acts as an omnidirectional barrier between these microorganisms and the body as well as a mediator of the immune response against attacks (2,3). To this end, a barrier of mucus, a viscoelastic gel, lines the colon and protects us from pathogens and harmful substances (4). The mucus is produced by goblet cells, which are specialized cells located in crypts of the colonic epithelial layer. Within the human colon, there are two mucus layers, of which the inner layer is attached to the colonic surface and the outer layer is unattached and is repeatedly removed from the colon using distal transportation to ensure the colonic lumen and wall remain healthy (5-7).

Within the outer mucus layer of the colon there lives a second population of microbiota named the mucus associated microbiota (MAM). The MAM differs from the LM in that they reside within the mucus layer, separated from the lumen. These bacteria use the glycans of mucins as a food source and assist in the digestion of otherwise indigestible carbohydrates, turning them into digestible short-chain fatty acids (SCFA), which are believed to play an important role in maintenance of gut and metabolic health (8). The MAM also stimulates the production of mucus and antimicrobial compounds against pathogenic bacteria (9-11). Furthermore, the MAM competes with pathogens for nutrients and produces metabolites to communicate with the host (12-14).

Indeed, disruption of the gut homeostasis is accompanied by several major health risks such as diabetes mellitus (T2D), inflammatory bowel disease (IBD), irritable bowel syndrome (IBS), and colorectal cancer (15,16). The colon is therefore intensively studied for its eminent role in human health, though research is hindered by a lack of conventional experimental techniques (17). Although there are a variety of established methods, the colon is not easily accessible for direct experimentation and colonoscopy is an invasive assay that does not provide cellular clarity (18-20). Furthermore, sequencing results of faecal samples are not representative of the complete microbiome, since layers of bacteria attached to the surface, known as biofilms, are not found (21-23). Faecal approaches also do not provide information about spatial variations in the microbiota, since the discharge is formed by all segments of the gastro-intestinal tract (24). Moreover, faecal approaches do not elucidate on the mucus layer.

To alleviate the difficulty of research in the gut, *in vitro* approaches have attempted to create accessible systems for gut research. Currently, these systems focus on the use of bioreactor (25) and organ-on-a chip (26,27) technology. An advanced dynamic continuous fermentation model was developed within TNO in 1999 (28, 29). This model, called the TIM-2 – an acronym for The Intestinal Model-2 – is an *in vitro* computer-controlled model which can simulate the large intestine. Within the TIM-2, pre-digested substrates are mixed with human microbiota under set pH and temperature levels which are continuously monitored. Peristaltic movements of the TIM-2 allow the model to simulate the mixing and transport. The model’s unique feature, the dialysis membrane, allows for the absorption of water and metabolites thereby preventing metabolite accumulation and ensuring physiological microbiome dynamics. The system is fully anaerobic, meaning the TIM-2 can be used to study fermentation. Samples can be taken from the lumen and from the dialysate, or “absorbed” fluid. Additionally, these systems allow for the examination of drug-microbiota interactions. The TIM-2 has a short adaptation period of only 16 hours, ensuring the composition of microbiota corresponds to the original inoculum. However, TIM-2 did not reproduce the colonic mucus layer, and thus is limited to lumen microbiota studies. Therefore, we aimed to incorporate a mucus compartment into the TIM-2 model (TIM-2*muc*) that mimics the functional response characteristics of the *in vivo* mucus layer when subjected to external stimuli. This innovation enables the simultaneous study of both the LM and the MAM, which has been challenging to achieve in both humans and available in vitro platforms.

To validate if the integration of the mucus compartment behaves as the *in vivo* MAM, we investigated the bacterial composition, alpha- and beta-diversity by 16S rRNA sequencing, and response of the MAM relative to the LM of the TIM-2*muc* and TIM-2 model, and SCFA production via gas-chromatography (GC). Our findings were further benchmarked against existing literature on MAM-LM differences.

## Material and Methods

In our study, we first collected faecal samples from eight healthy lean adult volunteers, each of whom was selected following stringent health criteria that precluded the use of probiotics, prebiotics, antibiotics, or any medications known to affect the gut environment. Volunteers used standardized collection kits designed to maintain anaerobic conditions during collection. After collection, individual samples were pooled to ensure standardization and reproducibility for subsequent experiments. Details about preparation of the pooled aliquots, the feeding medium and dialysate were described earlier (30).

The experimental phase of our study made use of advanced *in vitro* colon models, specifically the TIM-2 and the upgraded TIM-2*muc* models. The TIM-2*muc* model, hereafter also referred to as advanced model, is different from the TIM-2 model, or conventional model, in that it has a second compartment joined to the luminal compartment (**Fig. 1**).

**Fig. 1.**
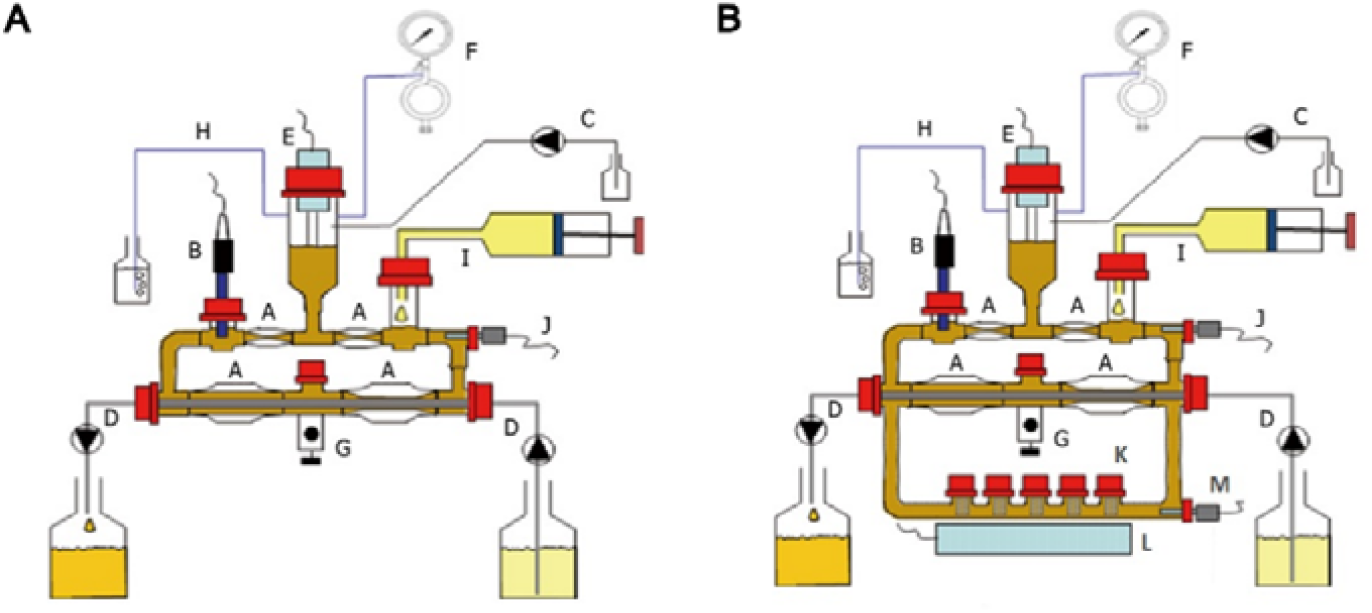
Side-by-side schematic overview of TIM-2 and TIM-2*muc*. **A** Schematic representation of the components of the conventional TIM-2 model. **B** Schematic representation of the components of the TIM-2*muc* model. Schematic components: A = Sleeves which mimic peristaltic movements, B = pH sensor, C = NaOH pump and inlet, D = Dialysis system which absorbs metabolites, E = Fluid level sensor, F = N_2_ pump and inlet, G = Luminal sample port, H = Bubbler to check for a closed gas circuit, I = Feed syringe, J = Luminal temperature sensor, K = Mucus compartment with five glass chambers, each with a metal cage containing mucus beads (Gut^3^Beads) and an electrospun polycaprolactone sheet coated with mucus (Gut^3^Gel), L = Temperature regulator, M = Mucus compartment temperature sensor

This mucus compartment was integrated into the setup as a series of glass chambers containing specially designed basket sinker (Basket Sinker, SS, Japanese Pharmacopeia, Univ.). Each sinker housed an electrospun polycaprolactone sheet (supplied by Vivolta, The Netherlands), which served as a scaffold representing the MAM. Prior to use, these sheets were sterilized by 70% ethanol and exposed to ultraviolet light, treated with sterile NaOH, and washed thoroughly with sterile phosphate-buffered saline. After drying, they were coated with colonic mucus gel (Gut^3^Gel by Bac^3^Gel, Portugal) and incubated overnight, for optimal adhesion conditions. The scaffold was then placed in the cages and surrounded by growth booster beads for colonic microbiota (Gut^3^Beads, Bac^3^Gel, Portugal), replicating the dual-layered mucosal structure observed *in vivo*. At the start of the experiments, the models were inoculated with faecal material which was allowed to adapt for 16 hours. As the luminal content further circulated through the system over a continuous period of 72 hours, it passed through the mucus compartment, allowing bacteria to attach to the scaffold and interact with the mucus in a manner similar to the naturally occurring MAM and LM interplay *in vivo*. Throughout the experiments, the colon models operated under controlled conditions, with computer-regulated adjustments to pH (5.8), temperature (37°C), and dialysis flow which ensured highly dense active and viable microbiota. Additionally, Gut^3^Beads were refilled every 24 hours.

Samples were collected every 24 hours (0-, 24-, 48-, and 72-hours) post-adaptation from both the lumen and the dialysis fluid. In the advanced TIM-2*muc* model, additional MAM samples were collected by extracting bacteria attached to the scaffold. This dual-sampling strategy enabled us to obtain a comprehensive profile of both LM and MAM communities.

Metabolic profiling was conducted using GC to quantify SCFA concentrations, with every measurement undergoing calibration and quality control. Complementary to these metabolic assessments, the microbial community was characterized using 16S rRNA gene sequencing (performed by Zymo Research, Germany), which were utilized in composition and diversity analyses.

## Results

To evaluate how the presence of mucus influences microbial dynamics, we assessed the microbiota composition by 16S sequencing analysis. The results revealed differences in the relative abundance of dominant phyla between the MAM and the LM. For instance, *Bacteroidetes, Actinobacteria*, and *Proteobacteria* were more abundant in the MAM compared to the LM throughout the experiment (**Fig. 2**).

**Fig. 2.**
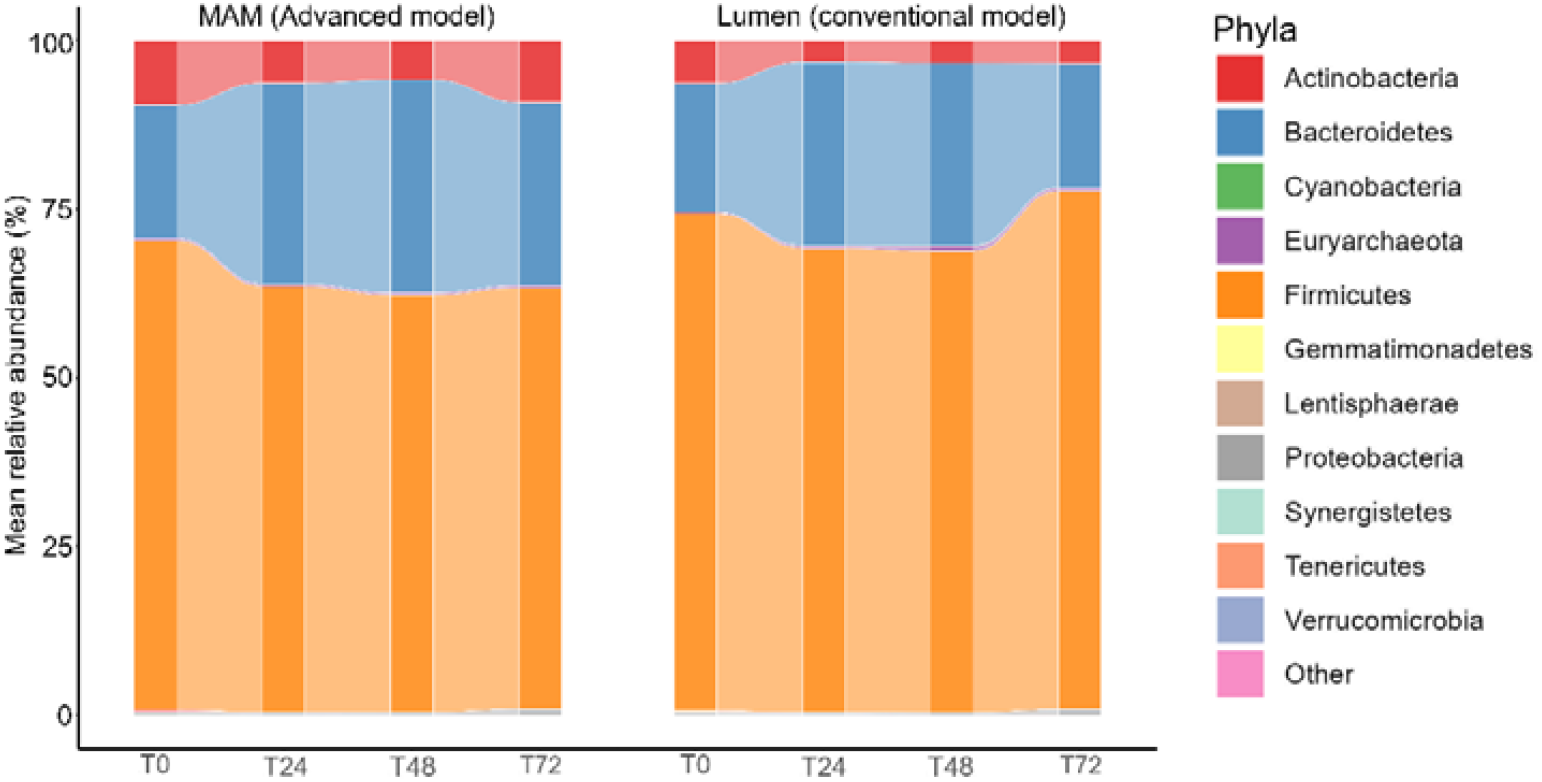
Comparison of the advanced (TIM-2*muc*) and conventional TIM-2 model at 0, 24, 48, and 72 hours of simulation (N=2). Alluvial stacked bar chart of microbiome composition at phyla level of MAM (advanced model) and Lumen (conventional model)

Whilst *Firmicutes* were more abundant in the LM than in the MAM. These data indicate that the microbial community composition in the MAM is distinct from that observed in the conventional LM. Unweighted Unifrac distances were calculated and visualized via Principal Coordinate Analysis (PCoA) to assess beta-diversity of microbiota samples derived from the advanced model (**Fig. 3A**). In the TIM-2*muc* system, the LM and MAM clustered closely together at T0, indicating early cross-compartment interactions. Over time, their trajectories diverged in a time-dependent manner, with distinct community shifts influenced by the presence of the mucus. Notably, the MAM and LM from TIM-2*muc* partially overlapped at T24 and T72 but were distinct at T48, suggesting that although these compartments harbor separate microbial communities, their trajectories ultimately converge. Rarefaction curves of the Chao1 index (**Fig. 3B**) revealed that overall alpha diversity declined over time. At T0 and T24, the diversity levels in the MAM and LM compartments were comparable (two-tailed t-test, p= 0.6752 and p = 0.1377, respectively), whereas at T48 and T72 the MAM exhibited significantly higher diversity (two-tailed t-test, p = 1.506e-10 and p = 1.071e-06), suggesting a greater capacity for species retention.

**Fig. 3.**
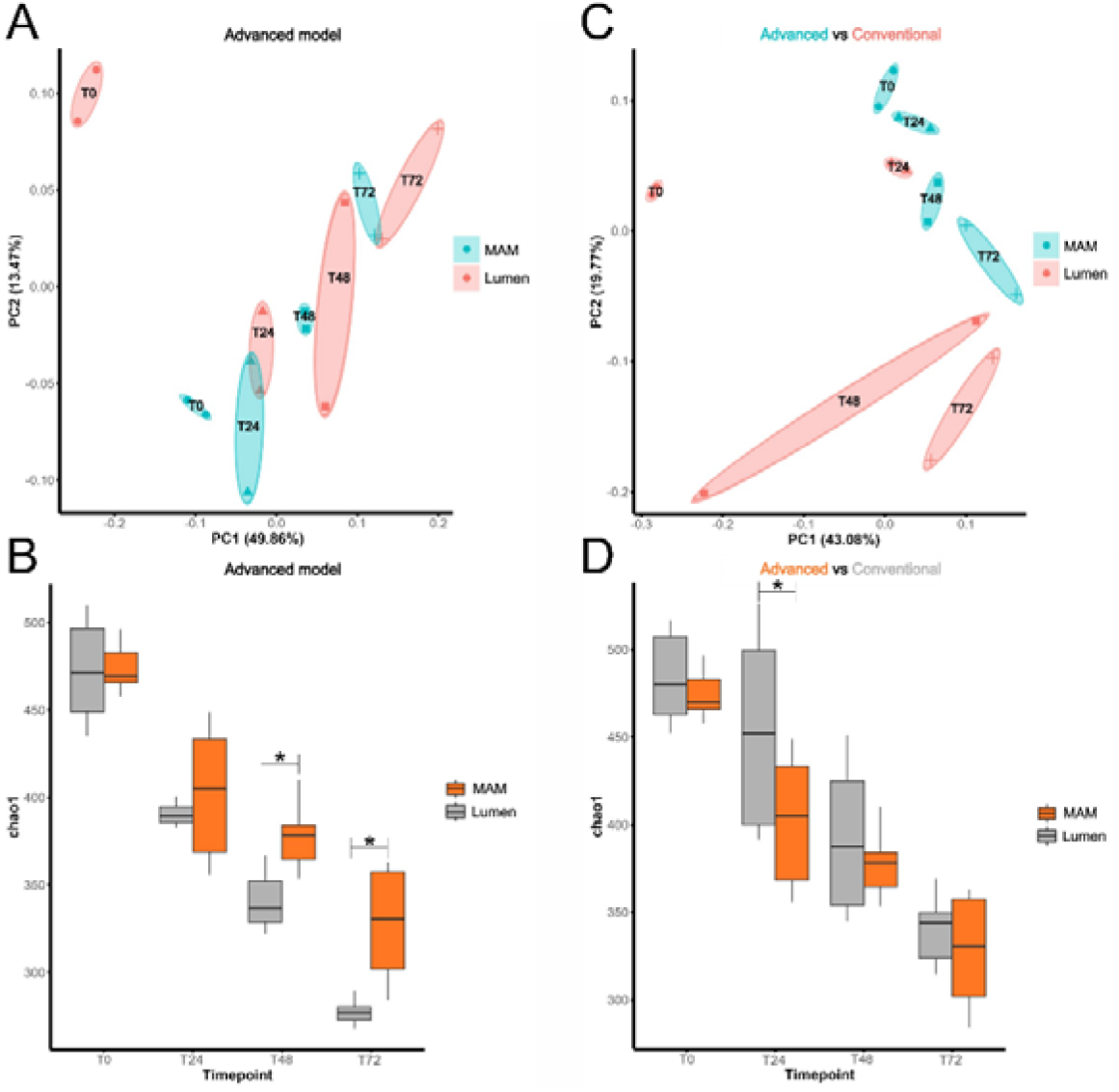
Comparison of the advan ed and conventional TIM-2 model at 0, 24, 48, and 72 hours of simulation) (N=2). Beta-and Alpha-diversity of advanced and conventional model compartments. (A,C) Unweighted Unifrac distances of MAM and lumen samples shown in PCoA plot. (B,D) Rarefaction Chao1 index of MAM and lumen samples shown in bo plots (^*^= significance).

In contrast, analysis of the conventional LM and MAM through Unweighted Unifrac distances and PCoA (**Fig. 3C**) showed distinct sample clustering, with no partial overlap between the MAM and LM. This distinct separation indicates that the introduction of mucus influences the composition of the microbiota. Furthermore, the MAM varied less over time compared to the LM, implying that it may provide a more stable environment and consequently microbial community due to bacterial attachment. Finally, rarefaction analysis of the Chao1 index for the conventional model LM and MAM (**Fig. 3D**) showed that, on average, the MAM had lower species richness than the LM. While differences at T0, T48, and T72 were not statistically significant (p = 0.0537, p = 0.0805, and p = 0.3978, respectively), a significant difference was observed at T24 (p = 0.0492). Collectively, these findings suggest that the introduction of mucus has an impact on the overall composition of the microbiome.

The production of SCFA was evaluated in both the advanced and conventional models using GC (**Fig. 4A**). Overall, the advanced model produced slightly lower total SCFA levels than the conventional model. However, the SCFA ratio consistently differed between the models at every time point (**Fig. 4B**). In the advanced model, butyrate accounted for a higher proportion of the total SCFA compared to the conventional model at T0, T24, T48, and T72, while acetate and propionate were present at lower ratios. These findings indicate that the inclusion of the mucus shifts the model’s metabolic profile toward enhanced butyrate production.

**Fig. 4.**
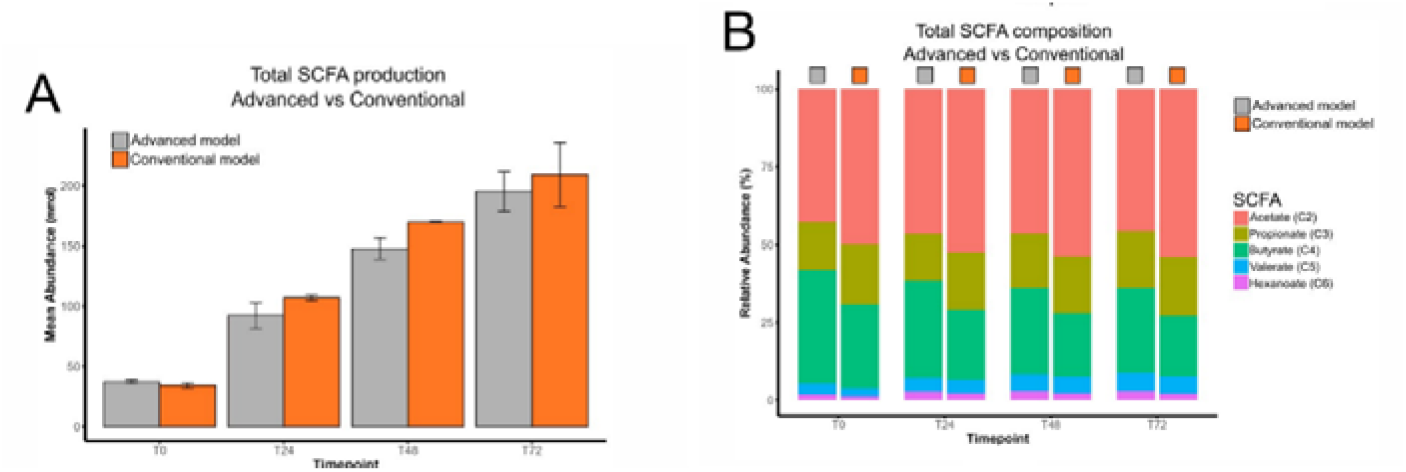
Comparison of the advanced and conventional TIM-2 model at 0, 24, 48, and 72 hours of simulation (N=2). Total SCFA production of the advanced and conventional model (A) and relative abundances of individual SCFA per timepoint (B).

The presented data demonstrates that incorporating mucus into the model fundamentally alters microbial community dynamics (composition and metabolic profiles). Most notable is the important role of mucus in stabilizing the luminal microbiota and enhancing butyrate production.

## Discussion

In this study, we investigated the TIM-2*muc* model, a novel adaptation of the conventional TIM-2 model that incorporates a mucus interface to simulate the MAM. The inclusion of this mucus matrix reshaped microbial dynamics. Furthermore, a notable increase in abundance of colonic mucus-associated genera was found in the MAM.

This study introduced a novel *in vitro* model for studying the MAM. In comparing the MAM to the conventional lumen, we observed increases in *Actinobacteria, Proteobacteria*, and *Bacteroidetes*, alongside a decrease in *Firmicutes*. These results largely concur with previous studies comparing MAM and luminal microbiota (31-32), except that our model showed a higher abundance of *Bacteroidetes*, contrary to earlier findings. However, several *Bacteroidetes* lineages possess mucin-degrading enzymes, giving them a competitive edge in the mucus layer and explaining their prominence in TIM-2*muc* (33). *In vivo*, oxygen leaking from the epithelium into the MAM creates a favourable environment for aerotolerant taxa and keeps strict anaerobes in check (32). Because TIM-2*muc* lacks this gradient, obligate anaerobes such as *Bacteroides* face no such constraint and can proliferate unhindered (34). Incorporating a controlled oxygen-diffusion step should therefore bring the model’s community structure closer to that of the native MAM.

Analysis of alpha- and beta-diversity revealed that introducing mucus reshaped the entire community. The MAM partly clustered with the advanced LM in PCoA, whereas the conventional LM formed a distinct cluster from the MAM, confirming that mucus alters luminal composition. Moreover, alpha-diversity was higher in the MAM than in the advanced LM but slightly below that of the conventional LM. This pattern mirrors *in vivo* brush-versus-stool work: brush-derived MAM communities differ from home-collected stool in beta-diversity, and stools typically display similar or higher alpha-diversity (35), mirroring the MAM to conventional LM contrast observed in our experiments; meaning the conventional LM resembles home-collected stool.

Another study comparing sampling methods reported that endoscopically collected stool, unlike home-collected stool, showed lower alpha-diversity and partial overlap with the MAM in unweighted UniFrac space (36). The advanced LM follows the same trajectory, its diversity sits below the MAM and partially overlaps with the MAM, suggesting that adding the mucus compartment nudged the luminal community from the home-collected stool profile, toward the profile of endoscopically collected stool.

Comparing SCFA production between TIM-2*muc* and TIM-2 revealed a lower total production and a shift in composition with addition of mucus. This is likely because the larger volume in TIM-2*muc* further dilutes the same amount of circulating feed and, consequently, total SCFA output falls. Moreover, the introduction of the mucus matrix creates a niche that draws microbial activity from the lumen, reducing its alpha diversity and thereby possibly the fermentative capacity. Although the mucus compartment contributes only a small biomass, its community favors butyrate production; a trait associated with mucosal bacteria (31). Since the LM is likely influenced by the MAM, it is logical to assume that the luminal environment would support increased butyrate producers as well.

Our study demonstrates that the TIM-2*muc* model represents an accurate and standardized approach to studying the MAM. By incorporating a dedicated mucus interface, the model not only dynamically reshapes the luminal environment to more closely mirror the *in vivo* state but also successfully recreates key features of the MAM observed in human studies. The advanced model showed an increased abundance of mucus-associated bacteria and distinct alpha- and beta-diversity patterns that align well with previously reported *in vivo* data, underscoring its physiological relevance.

## Acknowledgments

Neeltje van der Veen, Nouschka Verboon, Ineke Kwakernaak, Mans Minekus, Oumaima Bennar, Tom Gorissen, Meenu Sharma, InnoGI team and Bac3Gel team. Daniela Pacheco acknowledges the support of the European Innovation Council Accelerator programme (HORIZON-EIC-2023-ACCELERATOROPEN-01 ID: 190135075), which partially funded the customisation process of the mucus models to accommodate dynamic physiologic conditions.

## Author Contributions

SR performed all the experiments, analysed the results and wrote the manuscript. KL and MC helped design all aspects of the study, supervised and analyzed the results. OH, JL and AH contributed to develop TIM-2*muc* model. DP customized Gut3Beads for TIM-2*muc* model together with the InnoGI team. LV contributed to the SCFA experiment.

